# Complete Protection from SARS-CoV-2 Lung Infection in Mice Through Combined Intranasal Delivery of PIKfyve Kinase and TMPRSS2 Protease Inhibitors

**DOI:** 10.1101/2023.07.19.549731

**Authors:** Ravi Kant, Lauri Kareinen, Ravi Ojha, Tomas Strandin, Saber Hassan Saber, Angelina Lesnikova, Suvi Kuivanen, Tarja Sirnonen, Merja Joensuu, Olli Vapalahti, Tom Kirchhausen, Anja Kipar, Giuseppe Balistreri

## Abstract

Emerging variants of concern of SARS-CoV-2 can significantly reduce the prophylactic and therapeutic efficacy of vaccines and neutralizing antibodies due to mutations in the viral genome. Targeting cell host factors required for infection provides a complementary strategy to overcome this problem since the host genome is less susceptible to variation during the life span of infection. The enzymatic activities of the endosomal PIKfyve phosphoinositide kinase and the serine protease TMPRSS2 are essential to meditate infection in two complementary viral entry pathways. Simultaneous inhibition in cultured cells of their enzymatic activities with the small molecule inhibitors apilimod dimesylate and nafamostat mesylate synergistically prevent viral entry and infection of native SARS-CoV-2 and vesicular stomatitis virus (VSV)-SARS-CoV-2 chimeras expressing the SARS-CoV-2 surface spike (S) protein and of variants of concern. We now report prophylactic prevention of lung infection in mice intranasally infected with SARS-CoV-2 beta by combined intranasal delivery of very low doses of apilimod dimesylate and nafamostat mesylate, in a formulation that is stable for over 3 months at room temperature. Administration of these drugs up to 6 hours post infection did not inhibit infection of the lungs but substantially reduced death of infected airway epithelial cells. The efficiency and simplicity of formulation of the drug combination suggests its suitability as prophylactic or therapeutic treatment against SARS-CoV-2 infection in households, point of care facilities, and under conditions where refrigeration would not be readily available.

## INTRODUCTION

To productively infect cells, SARS-CoV-2 must fuse its lipid envelope membrane with membranes of the host cell^1^. This fusion event reaches its maximal efficiency when the viral surface protein spike (S) is exposed to low pH^2^ and results in delivery of the viral RNA genome into the cytoplasm of the host cell where synthesis of viral proteins and genome replication occur^3^. To trigger fusion, the viral S must first bind to a cellular receptor, e.g. angiotensin-converting enzyme 2 (ACE2)^4^, and then be cleaved by cellular proteases such as transmembrane serine protease 2 (TMPRSS2)^4^ localized at the cell surface and early endosomes, or cathepsin-L and -B in late endosomes and lysosomes^5^. Thus, depending on the pH of the extracellular space and the plasma membrane availability of proteases, the fusion can occur at the cell surface or, following virus endocytosis, in the endosomal system^2, 5^.

It has been shown previously that SARS-CoV-2 infection can be blocked by serine protease inhibitors such as nafamostat mesylate in cells that express TMPRSS2 but not cathepsins (e.g. Calu-3 cells)^6^. In cells that instead express cathepsins but not TMPRSS2 (e.g. VeroE6 or A549 cells), infection depends on the delivery of endocytosed viruses to endo/lysosomes, a process that can be efficiently inhibited by drugs that interfere with endosome maturation and acidification such as Bafilomycin A1, chloroquine or ammonium chloride. Infection was efficiently blocked in cells devoid of TMPRSS2 by inhibiting PIKfyve phosphoinositide kinase, a host enzyme involved in early-to-late endosome maturation, with either apilimod dimesylate or Vacuolin^6, 7^. In cells that express both TMPRSS2 and cathepsins, full inhibition of infection was achieved by combined use of nafamostat mesylate and apilimod dimesylate ^6^. Unexpectedly, however, this antiviral activity was synergistic even in Calu-3 cells that did not respond to apilimod dimesylate alone^6^ through a mechanism that remains to be elucidated.

The rise of immune-resistant variants, the challenge of delivering vaccines in under-developed countries, and the fraction of the global population that does not want to or cannot be vaccinated highlights the importance for the development of alternative prophylactic and curative strategies. Current antiviral therapies that complement vaccine treatment against SARS-CoV-2 are based on systemic delivery (i.e. oral or intravenous administration) of agents that directedly target viral components, including neutralizing monoclonal antibodies^8^, small molecules that interfere with viral RNA synthesis (e.g. remdesivir^9^, molnupiravir^10^), or inhibitors of viral proteases (e.g. Paxlovid^11^). The use and efficacy of these treatments, however, are limited by several important factors, including the rise of drug-resistant viral mutants^12, 13^, achieving the required effective drug concentration at the site of infection (i.e. the respiratory tract), bioavailability, and potential harmful side effects.

Because of the significantly lower mutational rate of the host genome, targeting host rather than virus factors required for infection has the potential to limit the rise of drug-resistant virus mutants. Targeted delivery to the respiratory track instead of systemic treatment should also decrease the required dosages, particularly when the treatment is intended for short periods of times (e.g., a few days or weeks). thereby lowering the risk for potential harmful side effects. Here we report that combined intranasal delivery of nafamostat mesylate and apilimod dimesylate in relatively small amounts during brief periods displayed a strong synergistic antiviral activity towards acute SARS-CoV-2 infection of laboratory mice together with a significant protective effect against cell death in the lower airway epithelium of the infected mice. We further report a drug formulation whose antiviral properties remained stable for over three months at room temperature, particularly beneficial for transportation in households, point-of-care facilities, and areas with limited access to refrigeration.

## RESULTS

### Combined treatment with apilimod dimesylate and nafamostat mesylate prevents SARS-COV-2 Wuhan, Alpha, Beta, Delta, or Omicron infections in cell culture

We recently demonstrated a synergistic full infection inhibition of chimeric VSV-SARS-CoV-2 (vesicular stomatitis virus, VSV, where the attachment and fusion glycoprotein G was replaced by the S protein of SARS-CoV-2 Wuhan) or native SARS-CoV-2 Wuhan by combined use of the PIKfyve phosphoinositide kinase inhibitors apilimod dimesylate or Vacuolin with the TMPRSS2 serine protease inhibitor, camostat mesylate^6^. A similar synergism was also observed using apilimod dimesylate together with nafamostat mesylate, another inhibitor of TMPRSS2^6^.

Here we confirmed that combined treatment with apilimod dimesylate and nafamostat mesylate of VeroE6 cells expressing low levels of TMPRSS2 was essential to inhibit infection by native SARS-CoV-2 Wuhan and extended similar observations to Alpha, Beta, Delta and Omicron variants^14^ (Fig. 1A, B, VeroE6-TMPRSS2). As shown before by us^6^ and others^4^, separate use of these inhibitors failed to fully prevent infection, as expected for cells that rely on TMPRSS2 and cathepsins as supplementary proteases in the complementary infection entry pathways.

**Figure 1.**
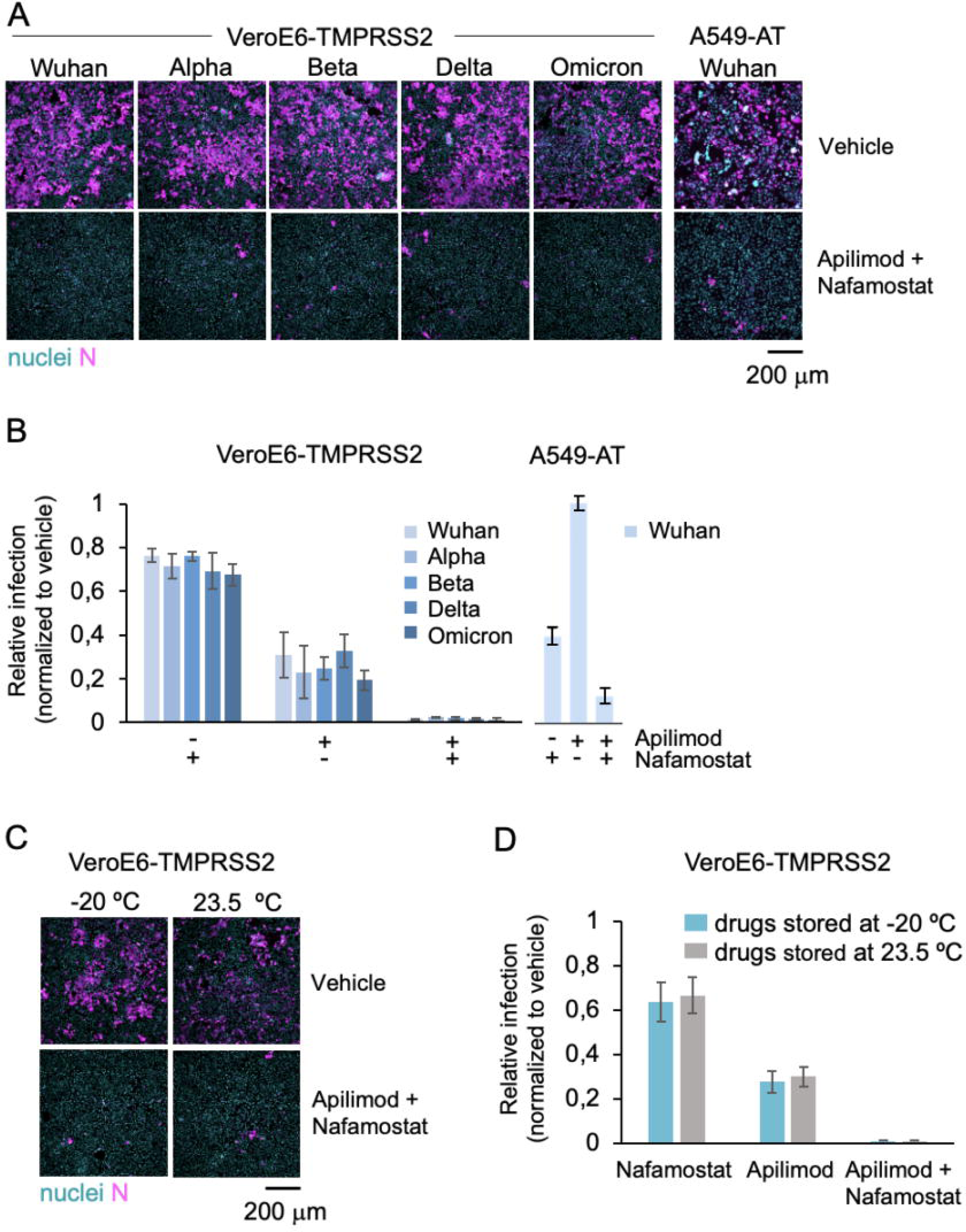
The combination of apilimod dimesylate and nafamostat mesylate inhibits SARS-COV-2 and its variants *in vitro*, and it is stable at room temperature. A. Representative fluorescence images of VeroE6-TMPRSS2 and A549-AT cells pre-treated with DMSO (Ctrl) or 2 μM apilimod dimesylate and 25 μM nafamostat mesylate, and one hour later infected for 20 h with indicated SARS-CoV-2 variants. Cells were stained with nuclear DNA dye Hoechst (nuclei, cyan) and immunostained with an antibody against the viral N protein (N, magenta). Scale bar = 200 μm. B-C. Quantification of the experiment shown in A. The percentage of N positive cells was determined by automated image analysis. Values represent the mean of three independent experiments and data are normalized to the infection levels obtained in DMSO vehicle treated infected cells (indicated as 1) in each experiment. The error bars represent the standard deviation. D. Representative fluorescence images of cells treated as in B with indicated drugs that had been stored either at -20 °C or room temperature (r.t.), for three months. One hour after drug treatment, cells where infected with SARS-CoV-2 Wuh strain for 20 h before fixation and immunofluorescence analysis as described in A. Scale bar 200 μm. E. Quantification of the experiment shown in D. The percentage of viral N positive cells was determined by automated image analysis. Values represent the mean of three independent experiments and data are normalized to the infection levels obtained in DMSO vehicle treated infected cells (indicated as 1) in each experiment. The error bars represent standard deviation.

Significant inhibition of SARS-CoV-2 Wuhan infection of human small lung adenocarcinoma derived A549-AT cells stably expressing ACE2 and high levels of TMPRSS2-eGFP was also obtained by combined use of apilimod dimesylate and nafamostat mesylate. (Figures 1 A, B; A549-AT).

### A stable aqueous formulation composed of apilimod dimesylate and nafamostat mesylate preserves its antiviral potency for several months

During our first experiments, we observed the development of turbidity of apilimod dimesylate and nafamostat mesylate, particularly at concentrations of 1 mg/ml or higher in solutions including PBS or DMEM culture media. Due to concerns about the potential decrease in effective drug concentration with aggregation or precipitation, particularly at the higher drug doses often required for animal studies, we investigated alternative formulations. We found that using water instead of salt-containing solutions as carriers for the drugs prevented precipitation; the solutions remained clear, even at concentrations of up to 1 mg/ml for apilimod dimesylate and 2 mg/ml for nafamostat mesylate when the mixture was dissolved in deionized water (Figures S1 A, B).

We then compared the antiviral potency of the stock solutions prepared in deionized water with the conventional way of preparing stock solutions dissolved in DMSO, or instead prepared in PBS or DMEM. We found that the antiviral potency in A549-AT cells of aqueous and DMSO stock solutions were indistinguishable from each other (Figures S2 A, B; Figure 1 B). Importantly, media containing drugs diluted from stocks made in PBS or DMEM and not water appeared toxic for cells, as they detached (Figures S2 C, D). Stock solutions directly prepared in deionized water were stable and maintain their antiviral activity, whether they were kept at -20 °C or 23.5 °C for up to 3 months (Figures 1 C, D). Based on these observations, all the animal inhibition infection studies that followed were undertaken using stock solutions of drugs dissolved in deionized water.

### Intranasal combined administration of apilimod dimesylate and nafamostat mesylate concurrent with virus inoculation prevents lung infection in mice

We used a non-lethal self-limiting mouse model of SARS-CoV-2 infection to test the antiviral effects of apilimod dimesylate and nafamostat mesylate, administered alone or in combination^15^. Wild type BALB/c mice were infected intranasally with a SARS-CoV-2 Beta isolate that harbours mutations (including N501Y) on the S protein known to enhance the interaction of the virus with the murine ACE2 receptor^16, 17^; this variant led to infection of the upper and lower airways and, to a lesser extent, of the pulmonary parenchyma (i.e. alveoli) 2 days after intranasal infection with a dose of 2 x 10^5^ plaque forming units (pfu)^15^.

A schematic timeline of the infection protocol is provided in Figure 2A. Mildly anesthetized mice received an intranasal dose of 5×10^5^ pfu of SARS-CoV-2 Beta suspended in 20 μl tissue culture media (DMEM containing 2% FBS), administered within 10-20 seconds after delivering 30 μl of either deionized water (vehicle control for the drugs) or apilimod dimesylate and nafamostat mesylate (diluted in deionized water), alone or in combination. Freshly dissolved drugs were administered intranasally twice daily, with 6-hour intervals, until the mice were euthanized for further analysis at 48 hours post-infection (hpi). The right lung of the mice was utilized for quantifying viral RNA levels using three primer sets by quantitative real-time PCR (qRT-PCR), targeting regions encompassing the viral RNA-dependent RNA polymerase (RdRp), and the envelope (E) protein gene (E, subE). The left lung and the head were used to evaluate pathological alterations and expression of viral nucleoprotein (NP) in upper airways and lungs by histology and immunohistology.

**Figure 2.**
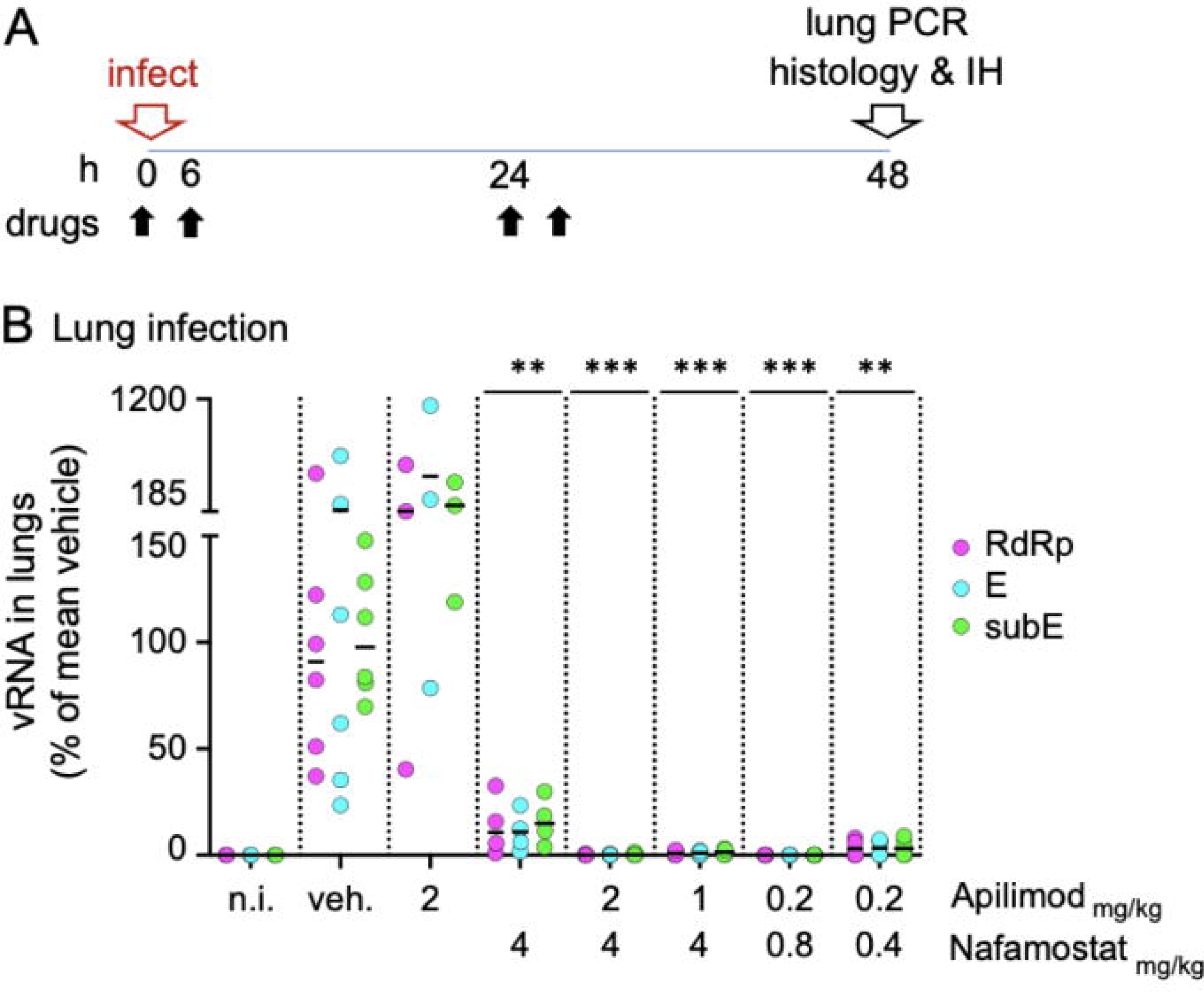
Intranasal delivery of combined drugs at low concentrations prevents SARS-CoV-2 beta lung infection in mice. A. Schematic description of the intranasal drug treatment and SARS-CoV-2 beta infection. Anesthetized mice received drugs intranasally in aqueous solution (30 μl) 10-20 seconds prior intranasal inoculation of SARS-CoV-2 beta (5×10^5^ plaque forming units in 20 μl DMEM). The drug treatment was repeated twice a day at 6 hours intervals at day 0 and day 1. At 48 hpi (day 2), mice were euthanized, and their right lung processed for real time quantitative PCR analysis to detect viral replication. The left lung and heads of the fixed animals were processed for immuno histology using anti N antibodies to monitor the integrity of the tissue and the distribution of viral antigens. B. PCR quantification of viral RNA in the lungs of mice treated with indicated drugs as described in A. For each mouse, the levels of viral RNA were detected using three non-overlapping primer sets, one targeting the viral genomic RNA dependent RNA polymerase gene (RdRp), and two sets against the viral gene E (E, subE). For each mouse, the obtained values were normalized first to the levels of actin in the same lung tissue and then to the mean viral RNA obtained in vehicle control treated infected mice (indicated as 100%). For each treatment, the mean (white bar) and standard deviation of the mean are indicated. The data were collected over two independent experiments each including vehicle controls. Each data point represents the RNA reads from one mouse. The concentration of apilimod dimesylate and nafamostat mesylate in mg/kg are indicated on the X axis. ***p<0.001.

A robust and reproducible infection was demonstrated in mice pre-treated with vehicle control by presence of viral RNA in the left lung (Figure 2 B, vehicle) and by expression of viral NP in sections of the contralateral lung, particularly in the respiratory epithelium in bronchioles and adjacent alveoli (type I and II pneumocytes); viral RNA (Fig 2 B) or viral NP were not detected in non infected control animals (Figure 3 A, non-infected and vehicle). Viral infection was associated with mild degeneration of respiratory and alveolar epithelial cells, together with some infiltrating neutrophils and mild peribronchial lymphocyte infiltration, features not observed in non-infected control mice (Supplementary Table 1). The nasal mucosa exhibited widespread viral NP expression in the respiratory and olfactory epithelium (Figure 3 B, vehicle; Supplementary Table 1).

**Figure 3.**
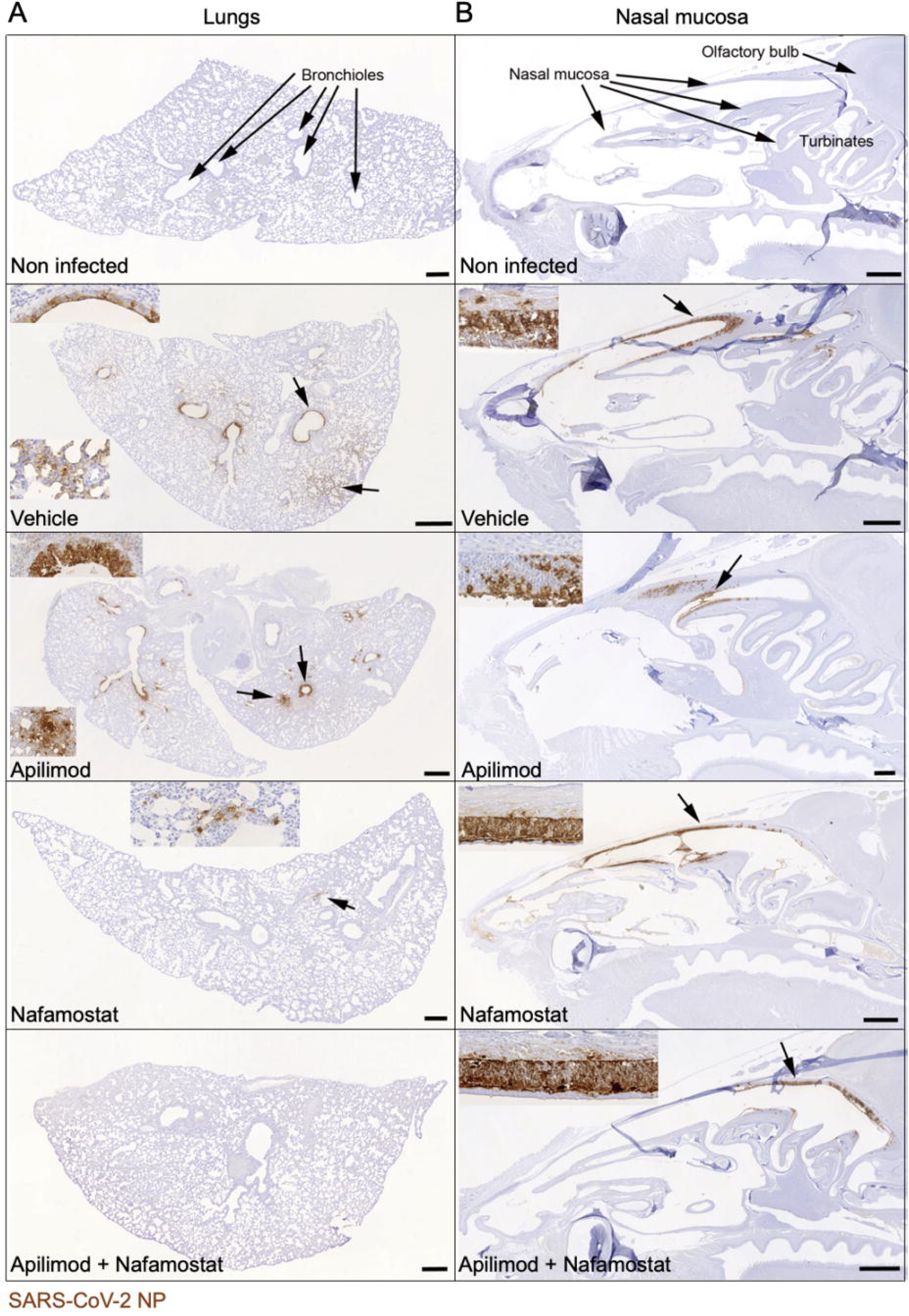
Immunohistological analysis confirms that intranasal apilimod dimesylate nafamostat mesylate treatment prevents lung infection and limits nasal infection. Immunohistology images of lungs and nasal mucosa from mice infected with the drugs as in Figure 2. Virus infected cells were identified with an antibody against the viral NP protein, using the horseradish peroxidase method (brown) and haematoxylin counterstain. Insets depict magnified images of the areas indicated by the arrows. Apilimod dimesylate: 2 mg/kg; nafamostat mesylate: 4 mg/kg; Apilimod dimesylate + nafamostat mesylate: 0.2 mg/kg + 0.8 mg/kg. Scale bars = 500 µm. A. Lungs. B. Nasal mucosa. Non infected mice exhibit no viral antigen in lung and nasal mucosa, whereas vehicle treated infected mice exhibit widespread SARS-CoV-2 NP expression in the lungs, both in bronchiolar epithelial cells and in pneumocytes in large groups of alveoli. This is also seen in epithelial cells in the entire nasal mucosa. In mice treated with Apilimod alone, the viral antigen expression pattern is identical, but its extent slightly reduced in the lung. After Nafamostat treatment, it is further reduced and only seen in small patches of alveolar epithelial cells. After combined Nafamostat and apilimod treatment, there is no evidence of viral antigen expression in the lung. In the nasal mucosa, positive cells are mainly seen in caudodorsal areas, in olfactory epithelial cells.

Intranasal treatment with apilimod dimesylate alone (2 mg/Kg) failed to prevent infection, as evidenced by presence of viral RNA in the lung and substantial expression of viral NP in nasal mucosa and lung (Figure 2B; Figures 3A-B). A similar lack of antiviral activity towards SARS-CoV-2 beta was reported for mice treated with apilimod dimesylate (50 mg/Kg daily) delivered intraperitoneally^18^.Intranasal treatment with nafamostat mesylate alone (4 mg/Kg) reduced lung infection as demonstrated by decrease in the amounts of viral RNA (Figure 2 B) and restricted viral NP expression in the lungs (Figure 3 A). The extent of nasal infection, however, appeared not to be affected by the treatment (Figure 3 B). A similar partial block of infection after intranasal delivery of SARS-CoV-2 Wuhan in animals treated intranasally with similar doses of nafamostat mesylate has been reported in Bablc mice transduced with adenoviruses expressing human ACE2 or in the highly susceptible transgenic K18-hACE2 mice expressing the human ACE2 under the control of the K18 promoter, highly expressed in the respiratory epithelial cells^18^, and hamsters that are also naturally infectable by the Wuhan strain of SARS-CoV-2^19^.

In glaring contrast to single drug treatment, combined intranasal administration of apilimod dimesylate and nafamostat mesylate fully prevented pulmonary infection over a wide range of concentrations even when using only 0.8 mg/Kg and 0.2 mg/Kg respectively. (Figures 2 B; Figure 3 A). Viral RNA levels were similar to or below detection limits (Figure 2 B), with no evidence of viral NP expression or tissue damage (Figure 3 A; Supplementary Table 1). While the combined drug administration regime did not prevent nasal infection as assessed by viral NP expression, it nevertheless appeared to be significantly less widespread and restricted to caudal areas of the respiratory and olfactory epithelium (Figure 3 B; Supplementary Table 1).

Our *in vivo* animal results are in full agreement with previous observations we obtained *in vitro* by infection of cells in culture conditions^6^, which showed strong inhibitory synergy rather than an additive inhibitory response to the combined delivery of both drugs. A quantitative assessment of the *in vivo* synergy is shown here by the enhanced decrease of viral RNA in lungs of mice treated with both drugs at very low concentrations (Figure 2 B, compare using 2 mg/Kg apilimod dimesylate and 4 mg/Kg nafamostat mesylate alone, and in combination).

### Intranasal combined delivery of apilimod dimesylate and nafamostat mesylate after infection strongly decreases bronchiolar cell death

We also tested the temporal span of protection offered by the combined intranasal delivery of either 2 mg/Kg apilimod dimesylate/4 mg/Kg nafamostat mesylate at 3 h.p.i., or 0.8 mg/Kg apilimod dimesylate/0.2 mg/Kg nafamostat mesylate at 6 hpi. Both regimes, compared to drug administration at the time of virus inoculation were equally less effective in diminishing the viral RNA load determined at 48 h.p.i. (Figure 4A), when (Figure 2 B; Figure 3 A). This lack of viral RNA load inhibition was expected for drugs administered 3-6 hours post virus inoculation, since at this time the first round of virus cell entry and concomitant onset of infection had already ensued.

**Figure 4.**
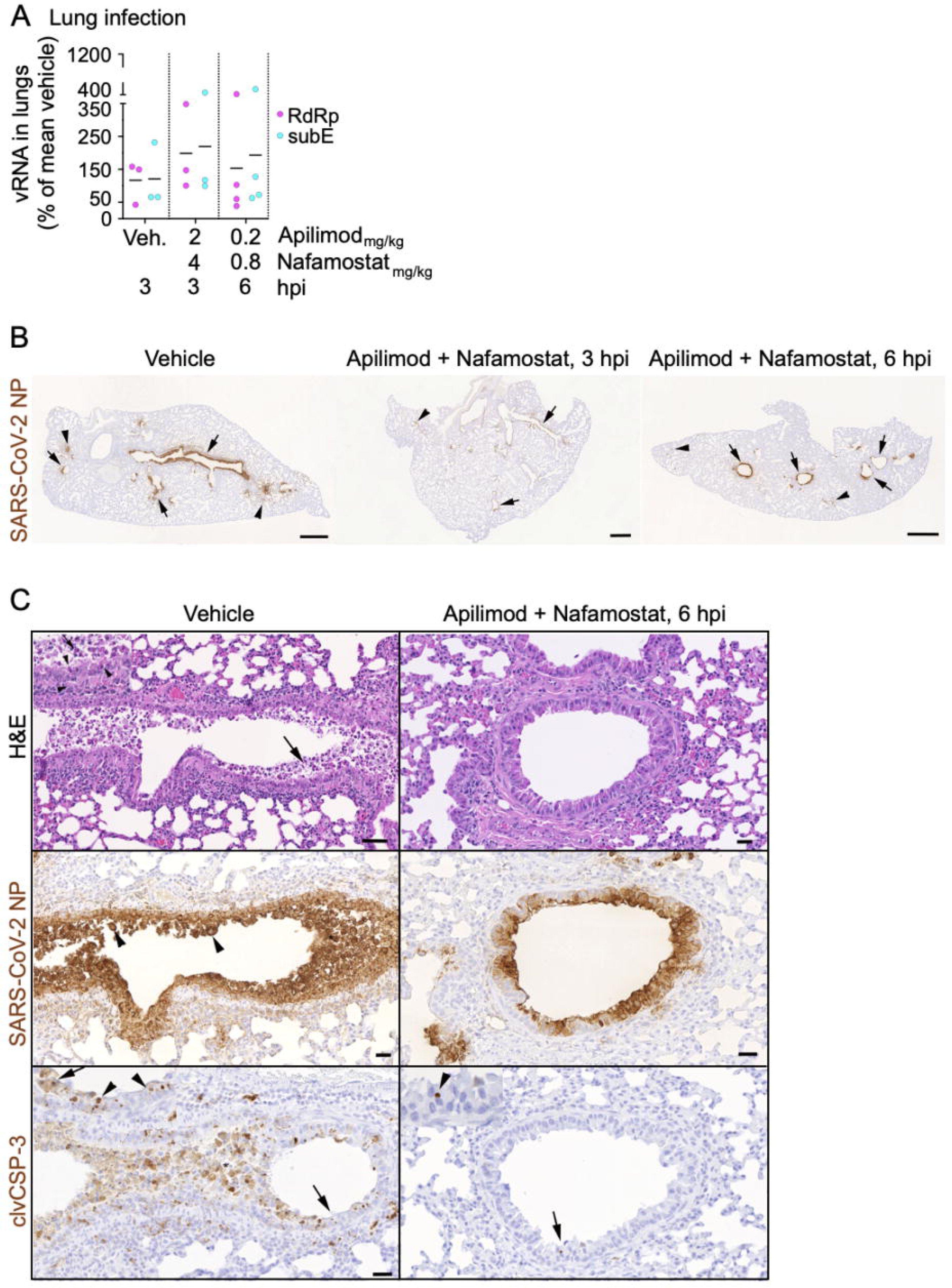
Intranasal delivery of combined drugs 3 h and 6 h post infection strongly decreases pulmonary cell death. A. PCR quantification of viral RNA in the lungs of mice treated intranasally with indicated drugs at 0 h (i.e., 10-20 seconds prior infection), 3 h and 6 h post SARS-CoV-2 beta infection. For each mouse, the levels of viral RNA were detected using two primer sets, targeting the viral genomic RNA dependent RNA polymerase gene (RdRp) and the sub-genomic viral gene E (subE), respectively. For each mouse, the obtained values were normalized first to the levels of actin in the same lung tissue and then to the mean viral RNA value obtained in vehicle control-treated infected mice (indicated as 100%). For each treatment, the mean (white bar) and standard deviation of the mean are indicated. Each data point represents the RNA reads from one mouse. The concentration of apilimod dimesylate and nafamostat mesylate in mg/kg, and the time of drug administration are indicated on the X axis. B, C. Immunohistology images of lungs from mice infected and treated as described in A. Virus infected cells were identified with an antibody against the viral N protein (B, C) and apoptotic cells were visualised with an antibody against cleaved caspase 3 (C), using the horseradish peroxidase method (brown) and haematoxylin counterstain. B. Mice treated with vehicle or with nafamostat (4 mg/kg) and apilimod (2 mg/kg), starting at 3 h.p.i. or 6 h.p.i. Vehicle treated mice exhibit widespread SARS-CoV-2 NP expression in epithelial cells of bronchi (arrows) and in groups of alveoli (arrowheads). With onset of treatment at 3 h.p.i., lung infection is seen, but is less widespread than in the vehicle treated animals. With onset of treatment at 6 h.p.i., viral antigen expression is also less extensive than in the vehicle treated animals, but the reduction is less marked. Scale bars = 500 µm. C. Mice treated with vehicle or with nafamostat (4 mg/kg) and apilimod (2 mg/kg), starting at 6 h.p.i. Consecutive sections of a bronchiole stained with hematoxylin-eosin (HE), for SARS-CoV-2 NP and for cleaved caspase-3. Insets represent higher magnifications of areas indicated by the arrows in the overview images. In the vehicle treated mouse, abundant degenerate cells are present in the lumen of the bronchiole, the epithelium exhibits several degenerating cells (arrowheads in inset). There is extensive viral NP expression in epithelial cells, including degenerate cells in the lumen (arrowheads). Staining for cleaved caspase-3 shows that infected epithelial cells die via apoptosis. Inset: apoptotic epithelial cells (arrowheads); arrow: sloughed off apoptotic epithelial cell). In the nafamostat and apilimod treated animals, the bronchiolar lumina are free of degenerate cells and the epithelium appears intact, although the majority of cells are virus infected as shown by the expression of viral NP. The extreme rarity of cleaved caspase 3-positive apoptotic cells (arrow; inset: arrowhead) confirms that infected cells are viable. Scale bars = 25 μm

As previously shown with vehicle-treated mice ^15^, infection of the bronchiolar epithelium was associated with substantial degeneration and sloughing of infected epithelial cells, as shown in consecutive sections stained with hematoxylin and eosin (HE) and for viral NP antigen (Figure 4 B). Staining of a further consecutive section for cleaved caspase-3 (clvCSP-3), a marker of apoptosis, confirmed that apoptosis was the main mode of cell death in the infected respiratory epithelium (Figure 4 B). The same staining approach was taken on the lungs of animals that had received apilimod dimesylate and nafamostat mesylate 3-6 h post virus inoculation, since the histology (HE stained section) suggested that there was less cell degeneration in the infected bronchioles. Indeed, viral NP expression was not associated with overt sloughing of infected epithelial cells, and there were only very rare apoptotic cells (Figure 4B).

These results show reduced cytopathic effect of the virus in the respiratory epithelium even when combined treatment with apilimod dimesylate and nafamostat mesylate was initiated after establishment of lung infection. .

### Intranasal combined delivery of apilimod dimesylate and nafamostat mesylate limits infection rebound

Lung infection determined 48 h.p.i. was prevented by treatment of mice with combined intranasal administration with 2 mg/Kg apilimod dimesylate and 4 mg/Kg nafamostat mesylate at the time of virus inoculation followed by a second dose 6 h.p.i. . To determine if infection rebound could occur when the drug treatment was stopped, we treated animals with vehicle or combined 2 mg/Kg apilimod dimesylate and 4 mg/Kg nafamostat mesylate twice daily for two days, and determined the levels of viral RNA and the extent of viral antigen expression in the lungs immediately after (at 48 h.p.i.) and two days later (at 96 h.i.) (Figure 5 A). The results confirmed that the combined drug treatment blocked lung infection over the 2-day treatment period (Figures 5 B, C). At 4 dpi, the vehicle-treated infected animals had not cleared the infection as their olfactory epithelium still harboured a few viral NP positive cells, consistent with limited residual infection (Supplemental Table 1). However, while there was evidence of lung infection based on the sporadic individual viral NP positive alveolar and bronchiolar cells (Figure 5 C; Supplemental Table 1), this infection was limited, as indicated by the lack of viral RNA in the lungs (Figure 5 B).

**Figure 5.**
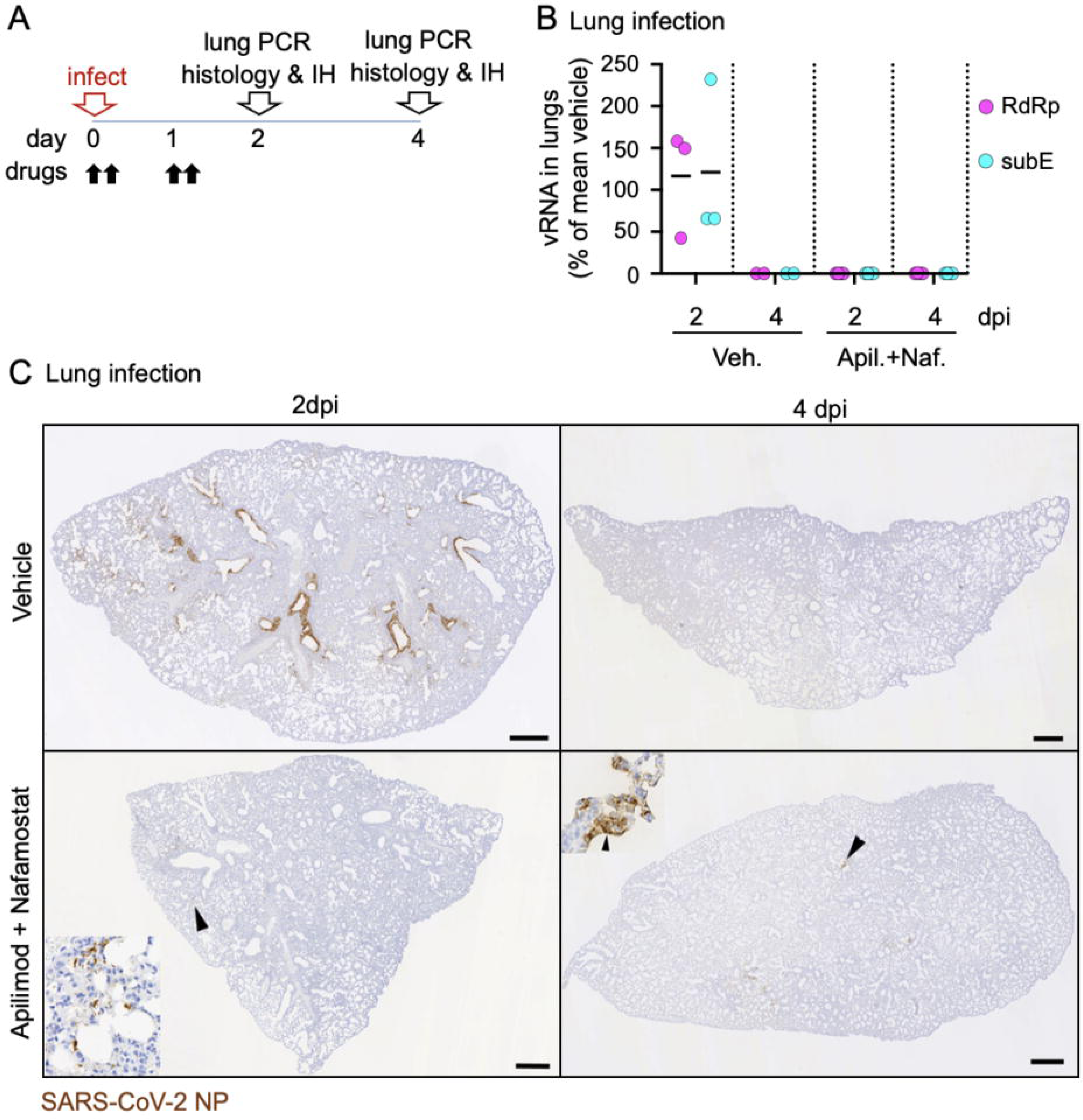
Intranasal delivery of combined drugs limits infection rebound. A. Schematic description of the intranasal drug treatment and SARS-CoV-2 beta infection in mice. The drug treatment was repeated twice a day at 6 hours intervals at day 0 and day 1. At 48 hpi (day 2) and 96 hpi (day 4) mice were euthanized and lungs processed for PCR analysis and Immunohystochemistry to detect viral RNA and the tissue distribution of infection, respectively. B. PCR quantification of viral RNA in the lungs of mice treated intranasally with indicated drugs and infected with SARS-CoV-2 beta. For each mouse, the levels of viral RNA were detected using two primer sets, one targeting the viral genomic RNA dependent RNA polymerase (RdRp) gene and other the sub-genomic viral gene E (subE). For each mouse, the obtained values were normalized first to the levels of actin in the same lung tissue and then to the mean viral RNA value obtained in vehicle control-treated infected mice (indicated as 100%). For each treatment, the mean (white bar) and standard deviation of the mean are indicated. Each data point represents the RNA reads from one mouse. For each treatment group, the day of euthanasia is indicated on the X axis. Apilimod dimesylate 2 mg/kg, nafamostat mesylate 4 mg/kg. C. Immunohistology for SARS-CoV-2 NP in the lungs at 2 and 4 days post infection with SARS-CoV-2 beta and treated as described above. Virus infected cells were identified with an antibody against the viral N protein, using the horseradish peroxidase method (brown) and haematoxylin counterstain. Bars = 500 µm. At 2 dpi, vehicle treated mice exhibit widespread SARS-CoV-2 N expression both in bronchiolar epithelial cells and in pneumocytes in groups of alveoli. In nafamostat and apilimod treated mice, there is no evidence of viral antigen expression. At 4 dpi, the infection has been cleared in the vehicle treated mouse, i.e. there is no evidence of viral antigen expression. In mice treated with nafamostat and apilimod for the first two days after intranasal virus challenge, there are a few small groups of alveoli with viral antigen expression in pneumocytes (inset: arrowhead; the inset is a higher magnification of the area highlighted by the arrowhead in the overview image).

## DISCUSSION

In this animal study we show potent block of SARS-CoV-2 infection by combined intranasal drug delivery of inhibitors for the host phosphoinositide kinase PIKfyve and TMPRSS2serine protease required for infection. Combined simultaneous intranasal delivery of both drugs was essential to prevent infection in mice inoculated intranasally with a human isolate of SARS-CoV-2 Beta, following the same inhibitory synergy we previously uncovered using native SARS-CoV-2 Wuhan and VSV-SARS-CoV-2 chimeras in *in vitro* tissue culture experiments ^6^, and repeated here with native SARS-CoV-2 Wuhan and Alpha, Beta, Delta and Omicron variants.

Robust inhibition of mice infection was obtained using drug concentrations significantly lower when compared to those reported in mouse COVID-19 models treated with other antivirals^11, 20^. For example, two consecutive doses of 0.8 and 0.2 mg/Kg delivered intranasally were sufficient in our study to block infection 2 d.p.i. In contrast, orally administered 300-1000 mg/Kg Paxlovid (viral RNA polymerase inhibitor)^11^ and 10-30 mg/Kg Remdesivir (viral protease inhibitor)^20^ administered twice daily to BALB/c mice was required for full lung protection determined 2 d. p.i. It remains to be determined whether the required differences in drug amounts are explained by the route of administration, e.g., intranasal vs oral.

We note that intranasal administration of these drugs, alone or in combination, did not induce clinical signs or histological changes in nasal cavity and lung that would suggest harmful effects in mice. Safety of Nafamostat or apilimod for humans, when used alone and administered intravenously or orally, has already been shown by the preliminary outcome from Phase I clinical trials. It remains to be determined whether their combined use, as proposed here, will also be tolerated by humans.

Importantly, here we show that aqueous solutions of apilimod dimesylate and nafamostat mesylate prepared with no other salts are stable and maintain inhibitory infection activity even after months of storage at room temperature. The enhanced solubility and long-term stability of stock aqueous solutions of nafamostat mesylate and apilimod dimesylate prepared with no other salts highlight the importance if these formulations when considering their potential use for therapeutics or prophylaxis against SARS-CoV-2 infection in households, point of care facilities, and in places where refrigeration would not be readily available.

Finally, we suggest the possibility of developing a system for intranasal delivery, using relatively low concentrations of the drugs, provided by a simple nasal spray or as an inhalation-administered treatment. The concept of access to an acute treatment is appealing, particularly in situations where a new SARS-CoV-2 variant might emerge that is unresponsive to previous immunization and for which a fast response is required before appropriate vaccines are available.

## MATERIAL AND METHODS

### Animals

A total of 30 female BALB/c mice (Envigo, Indianapolis, IN, USA) were transferred to the University of Helsinki biosafety level-3 (BSL-3) facility and acclimatized to individually ventilated biocontainment cages (ISOcage; Scanbur, Karl Sloanestran, Denmark) for seven days with ad libitum water and food (rodent pellets). For subsequent experimental infection, the mice were placed under isoflurane anaesthesia and inoculated intranasally with 20 µl of virus dilution or DMEM (non-infected control). The animals were held in an upright position for a few seconds to allow the liquid to flush downwards in the nasal cavity. All mice were weighed daily. Their wellbeing was further monitored carefully for signs of illness (e.g. changes in posture or behaviour, rough coat, apathy, ataxia). Euthanasia was performed under terminal isoflurane anaesthesia with cervical dislocation. Experimental procedures were approved by the Animal Experimental Board of Finland (license number ESAVI/28687/2020).

### Cells

A549 stably expressing human *ACE2* and *TMPRSS2* fused to GFP(A549-AT) were generated by transduction in two sequential steps. First, by using a third-generation lentivirus pLenti7.3 ACE2 blasticidine^14^ where the expression of blasticidine is driven by a separate promoter downstream of the *ACE2* coding sequence. Second, after selection of transduced cells with 3 μg/ml blasticidine for 7 days, the resulting A549-ACE2 cells where transduced again with a commercially available third-generation lentivirus encoding human *TMPRSS2* with a C-terminally fused histidine tag followed by the mGFP protein, and using a separate promoter to express the puromycin resistant gene (Angio-Proteomie, catalogue number vAP-0101). Lentiviral infections were carried in Dulbecco’s modified eagle’s medium (DMEM), 2 mM glutamine, 0,5 % bovine serum albumin (BSA), and 1x penicillin/streptomycin antibiotic mix, at an MOI of 0.3 infectious units/cells (lentivirus titre provided by the manufacturer’s). After 7 day selection with 3 μg/ml puromycin, the cell population was were expanded with three sequential passages and stored in liquid nitrogen. During infection experiments, blasticidine and puromycin were not included in the media. VeroE6-TMPRSS2^14^ and A549-ACE2-TMPRSS2-GFP^21^ were grown at 37 °C and 5% CO_2_ in DMEM supplemented with 2 mM glutamine, 10 % foetal bovine serum (FBS), and 1x penicillin/streptomycin antibiotic mix.

### Virus isolation, propagation, and sequencing

The Wuhan/D614G and Alpha, Beta, Delta, Omicron SARS-CoV-2 variants viruses were isolated from infected patient nasopharyngeal samples as described^14^ and amplified using transmembrane serine protease 2 (TMPRSS2)-expressing Vero E6 cells (VeroE6-TMPRSS2^14^). Once propagated, their genomic sequence was confirmed using an Illumina platform available at the Department of Virology, University of Helsinki^14^. The Wuhan and beta variant sequences are described in detail in^15^ and have been deposited in the NCBI GenBank database under accession numbers MZ962407 and MW717678, respectively. At three days post inoculation, the collected medium from the infected cell dishes was centrifuged twice at 4500xg for 10 min at 4°C, and the cleared supernatant aliquoted in cryotubes and stored at -80 °C in the BSL3 facility. All viruses were propagated in Minimum essential medium (MEM) containing 2% FBS, 20 mM HEPES, pH 7.2, 2 mM glutamine and 1x penicillin/streptomycin antibiotic mix. Virus titrations were performed by standard plaque assay in VeroE6-TMPRSS2 cells as previously described^14^. Infected cells were maintained in incubators at 37 °C and 5% CO_2_ in the BSL3 facility of the Helsinki University Hospital.

### Virus infection and drug treatments in cell cultures

VeroE6-TMPRSS2 and A549-ACE2-TMPRSS2-GFP cells were seeded in MEM containing 2% FBS, 20 mM HEPES pH 7.2, 2 mM glutamine and 1x penicillin/streptomycin antibiotic mix at 15,000 cells per well in 96 well imaging plates (catalogue number 6005182; PerkinElmer) 24 h before infection in the same medium with the various SARS-CoV-2 strains. Stocks of Apilimod dimesylate (Tocris, catalogue number 7283) and Nafamostat mesylate (Tocirs, catalogue number 3081) were diluted in the same medium and added at indicated times. The amount of virus used to infect the cells was adjusted to obtain15-20% infected cells as determined by immunofluorescence 20 h.p.i. using high-content imaging and automated image analysis after fixation with 4 % paraformaldehyde (in PBS), for 20 min at room temperature.

### Immunofluorescence

Fixed cells were washed three times with Dulbecco-modified PBS containing 0.2% BSA (DPBS/BSA), permeabilized with 0.1% Triton X-100 in DPBS/BSA and processed for immunodetection of viral N protein, automated fluorescence imaging, and image analysis. Briefly, viral NP was detected with an in-house-developed rabbit polyclonal antibody^22^ counterstained with Alexa Fluor 647-conjugated goat anti-rabbit secondary antibody (ThermoFisher Scientific, catalogue number A32733); nuclear staining was done using Hoechst DNA dye (ThermoFisher Scientific, catalogue number H3570). Automated fluorescence imaging was done using a Molecular Devices Image-Xpress Nano high-content epifluorescence microscope equipped with a 10× objective and a 4.7-megapixel CMOS camera (pixel size, 0.332 μm). Image analysis was performed with CellProfiler-4 software (www.cellprofiler.org). Automated detection of nuclei was performed using the Otsu algorithm inbuilt in the software. To automatically identify infected cells, an area surrounding each nucleus (5-pixel expansion of the nuclear area) was used to estimate the fluorescence intensity of the viral NP immunolabeled protein, using an intensity threshold such that <0.01% of ‘positive cells’ were detected in noninfected wells.

### Mice infection and drug treatments *in vivo*

9-week-old female BALB/c mice were anesthetized using isoflurane intranasally inoculated (n = 4 per group) with 2 × 10^5^ plaque forming units (PFU) in 20 μl of DMEM of the beta variant of SARS-CoV-2, or mock-infected with deionized water (n = 2)^15^. Drugs solubilized in deionized water were administered intranasally in 50 μl volume per mouse, at the indicated time points. The 50 μl drop was applied at the opening of the animal nostrils and the liquid was naturally inhaled while breathing. At 2 or 4 d.p.i., animals were euthanized under terminal isoflurane anaesthesia with cervical dislocation and dissected immediately after sacrifice; the right lungs were dissected and collected and frozen at -80 °C before PCR analysis of viral RNAs, whereas the left lung and heads were fixed in 10% buffered formalin for 48 h and stored in 70% ethanol for histological and immunohistochemical examinations.

### RNA Isolation and RT-qPCR

RNA was extracted from lung samples using Trizol (Thermo Scientific) according to the manufacturers’ instructions. Isolated RNA was subjected to one-step RTqPCR analysis as described using primers specific for the viral genome encoding for the RNA-dependent RNA polymerase (RdRp)^23^ and for E^24^ genes with TaqMan fast virus 1-step master mix (ThermoFisher Scientific, catalogue number 4444432) using AriaMx instrumentation (Agilent, Santa Clara, CA, USA). The actin RT-qPCR is described in^25^.

Primer and probe sequences used in the RT-qPCR.

Target Sequence

RdRp Forward gtgaratggtcatgtgtggcgg^23^

Probe caggtggaacctcatcaggagatgc^23^

Reverse caratgttaaasacactattagcata^23^

Subgenomic E Forward cgatctcttgtagatctgttctc ^24^

Probe acactagccatccttactgcgcttcg^24^

Reverse atattgcagcagtacgcacaca ^24^

Genomic E Forward acaggtacgttaatagttaatagcgt ^24^

Probe acac-tagccatccttactgcgcttcg ^24^

Revere atattgcagcagtacgcacaca ^24^

Beta-actin Forward actgccgcatcctcttcct ^25^

Probe cctggagaagagctatgagctgcctgatg ^25^

Reverse tcgttgccaatggtgatgac ^25^

### Histology and Immunohistochemistry

The left lungs and heads of the sacrificed mice were trimmed for histological examination and paraffin-wax embedded. The heads were sawn longitudinally in the midline using a diamond saw (Exakt 300; Exakt, Oklahoma, OK, USA), then decalcified and processed as previously described^15^. Consecutive sections (3–5 µm) were prepared from lungs and heads and stained with haematoxylin–eosin (HE) or subjected to immunohistology for the detection of SARS-CoV-2 antigen expression using a rabbit polyclonal anti-SARS-CoV NP antibody that cross reacts with NP of SARS-CoV-2 (Rockland Immunochemicals, Limerick, USA, catalogue number 200-402-A50); cleaved caspase 3 was detected with the antibody (rabbit anti-cleaved caspase-3 (Asp175), clone 5A1E; 9664; Cell Signalling Technologies) as previously described^15^.

---

**Supplementary figure 1.**
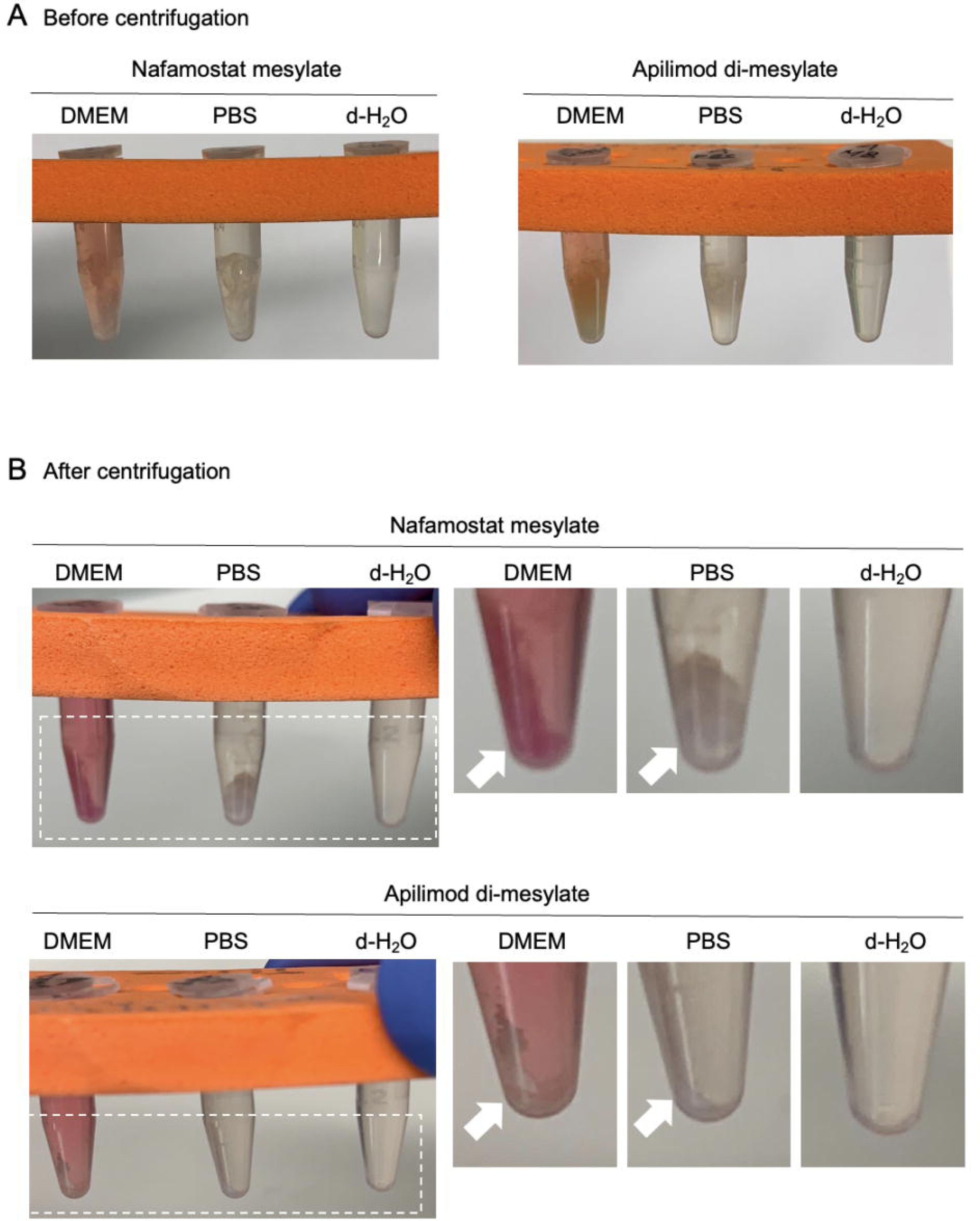
Solubility of apilimod dimesylate and nafamostat mesylate in different solutions. A. Apilimod dimesylate (1 mg/ml) and nafamostat mesylate (2 mg/ml) were solubilized in DMEM, PBS, or de-ionized water (d-H_2_O). A visible cloudy precipitate formed when the drugs were solubilized in DMEM and PBS, but not in d-H_2_O. B. After centrifugation at 10.000xg for 1 minute, at room temperature, a visible pellet formed if the drugs were solubilized in DMEM or PBS (white arrows), but not in the drugs solubilized as in A precipitate d-H_2_O.

**Supplementary figure 2.**
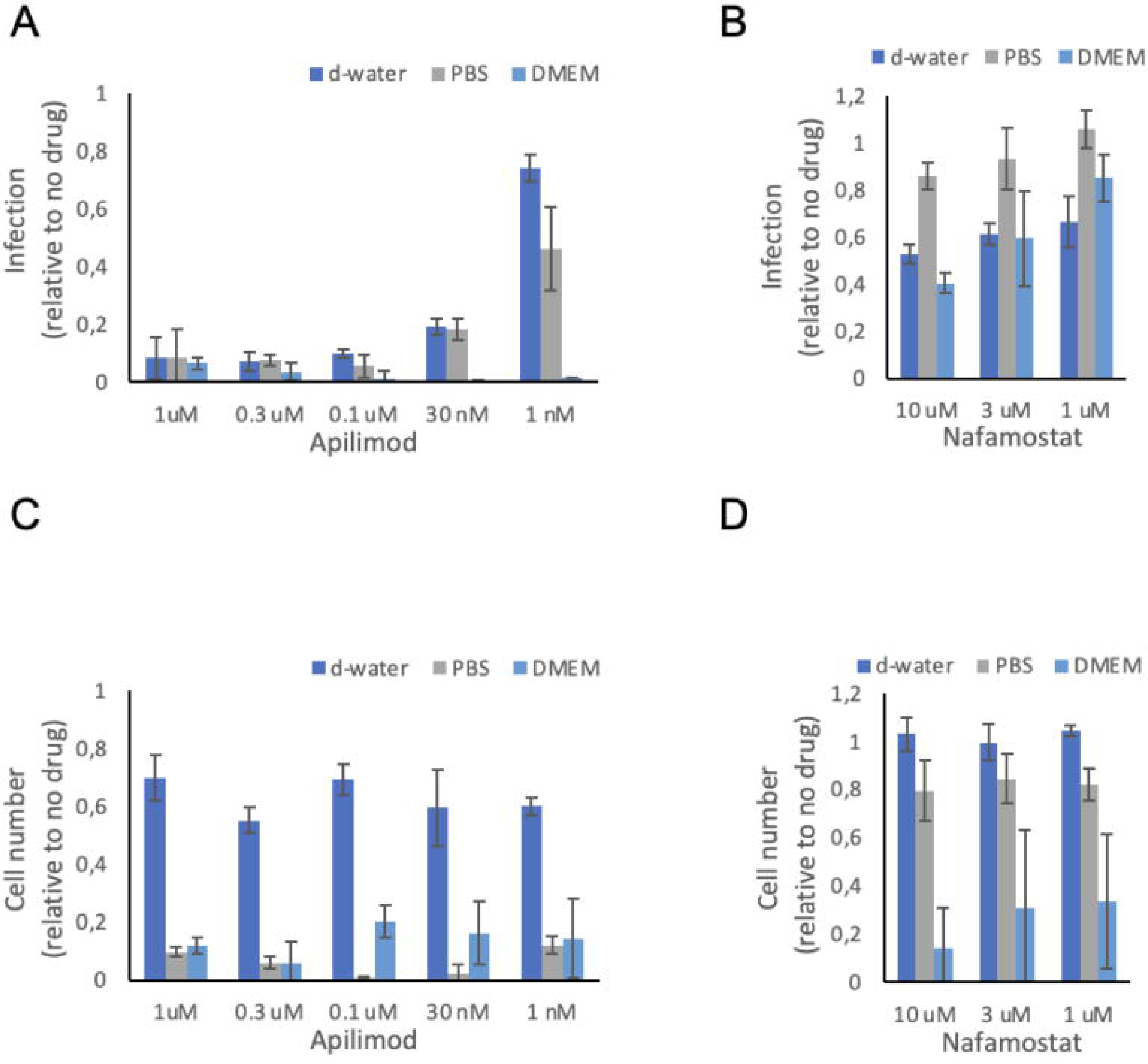
Antiviral activity and cell toxicity of apilimod dimesylate and nafamostat mesylate in different solutions. Stock solutions of apilimod dimesylate (1 mg/ml) and nafamostat mesylate (2 mg/ml) were prepared in DMEM, PBS, or de-ionized water (d-H_2_O). The drugs were further diluted in DMEM 2%FBS and added to Vero-E6 (apilimod dimesylate) or A549-AT cells (nafamostat mesylate) in 96-well plates at the indicated concentrations, 30 min before infection with SARS-CoV-2 Wuh (MOI= 0.5). Cells were fixed at 20 hours post infection and processed for immunofluorescence analysis using antibodies against the viral protein N to identify infected cells and Hoechst DNA staining to identify the cells nuclei. High-content imaging and automated image analysis were used to determine the number of infected cells in each sample. A-B. Number of infected cells for each drug treatment relative to the values obtained from infected cells treated with vehicle controls (indicated as 1). Values represent the mean and standard deviation of three independent experiments. C-D. Number of cells attached to the surface of the plates for each drug treatment relative to the values obtained from infected cells treated with vehicle controls (indicated as 1). Values represent the mean and standard deviation of three independent experiments.

## ACKNOWLEDGEMENTS

The authors are grateful to the team of laboratory technicians in the Histology Laboratory, Institute of Veterinary Pathology, Vetsuisse Faculty, University of Zurich, for excellent technical support. High-throughput imaging was performed at the Light Microscopy Unit of the University of Helsinki.

This research was supported by the Academy of Finland (grant numbers 335527 to G.B., 351040 and 336490 to O.P.V., 339510 to T.Si., 321809 to T.St.); European Union’s Horizon Europe Research and Innovation Program grant 101057553 (G.B., O.P.V.); Finnish Institute for Health and Welfare; VEO—European Union’s Horizon 2020 (grant number 874735 to O.P.V. and T.S.); Helsinki University Hospital funds TYH2021343 (O.P.V.); the Jane and Aatos Erkko Foundation (to O.P.V. and T.Si.); the University of Helsinki Graduate Program in Microbiology and Biotechnology (R.O.); Helsinki Institute for Life Sciences (HiLIFE, G.B. and O.P.V.), the Australian Research Council (ARC) Discovery Early Career Researcher Award (DE190100565 to M.J.) and The University of Queensland Amplify fellowship (to M.J). S.H.S. and N.Y. were supported by the Australian Government Research Training Program (RTP) and Commonwealth Tuition Fee Offset Scholarship, and N.Y. by The Westpac Scholars Trust. T.K. was supported by NIH Maximizing Investigators’ Research Award (MIRA) GM130386 and NIH Grant AI163019.

The funders had no role in the study design, data collection, or interpretation. We declare no competing interests.

## AUTHOR CONTRIBUTIONS

G.B. and T.K conceived the project. G.B. and A.K. planned and guided the work. R.K. and L.K. performed all the experiments in mice in BSL3. T.St. performed all the PCR analysis. R.O. and G.B performed the cell culture experiments. S.H.S., A.L. and S.K. produced virus stocks, titered the stocks, and helped with the PCR experiments and analysis. T.S., O.P.V., and M.J., coordinated the work in BSL3, provided resources, and funding. M.J. and S.H.S. helped with preparing the figures. G.B., A.K. and T.K. wrote the manuscript, analyzed and interpreted the results. All authors participated in proof-reading the manuscript and approved it for publication.

## Notes

### Competing Interest Statement

The authors have declared no competing interest.

