## Supplementary Table 1 for "Complete Protection from SARS-CoV-2 Lung Infection in Mice Through Combined Intranasal Delivery of PIKfyve Kinase and TMPRSS2 Protease Inhibitors": Supplementary Table 1 Animal data.docx

**Supplementary Table S1**. Histological changes, SARS-CoV-2 nucleoprotein and RNA expression in 9-week-old female BALB/c mice infected with 2 x 10^5 PFU SARS-CoV-2 Beta variant and euthanized at 2 or 4 days post infection and with different treatment regimes.

| **Treatment**  [Animal no] | **DPI** | **Histological changes (lung) and viral antigen expression (head*, trachea, lung, bronchial lymph node)** | **Virology (lung; PCR)** | | |
| --- | --- | --- | --- | --- | --- |
|  |  |  | **RdRp** | **subE** | **E** |
| **Experiment 1** | | | | | |
| **Not infected, not treated**  [1.1]  Fig 3A | **2** | **Lung (HE)**: NHA | 0 | 0 | 0 |
|  |  | **Nose**: neg  **Trachea**: neg  **Lung**: neg  **BLN**: NE |  |  |  |
| **Not infected, not treated**  [1.2]  Fig 3B | **2** | **Lung (HE)**: NHA | 0 | 0 | 0 |
|  |  | **Nose**: neg  **Trachea**: neg  **Lung**: neg  **BLN**: NE |  |  |  |
| **Infected, not treated**  [1.3]  Fig 3A | **2** | **Lung (HE)**: a few larger bronchioles with several degen EC; mild pb LC infiltration; one area with activated type II pc, some degen AEC and infiltrating NL | 99.31 | 113.03 | 147.77 |
|  |  | **Nose**: disseminated patches of pos REC and OEC  **Trachea**: some degen pos EC in lumen  **Lung**: many bronchioles with several individual to almost all pos EC; vAg covering luminal surface; adjacent large areas of alveoli with pos type I and II pc  **BLN**: several pos macrophages/DC |  |  |  |
| **Infected, not treated**  [2.1]  Fig 3B | **2** | **Lung (HE)**: a few larger bronchioles with several degen EC; mild pb LC infiltration; focal area with activated type II pc, some degen AEC and infiltrating NL | 82.36 | 35.27 | 80.85 |
|  |  | **Nose**: disseminated patches of pos REC and OEC  **Trachea**: some degen pos EC in lumen  **Lung**: many bronchioles with several individual to almost all pos EC; vAg covering luminal surface; adjacent large areas of alveoli with pos type I and II pc  **BLN**: several pos macrophages/DC |  |  |  |
| **Infected, not treated**  [2.2] | **2** | **Lung (HE)**: a few larger bronchioles with several degen EC; mild pb LC infiltration | 122.26 | 250.82 | 83.7 |
|  |  | **Nose**: disseminated patches of pos REC and OEC  **Trachea**: some degen pos EC in lumen  **Lung**: several bronchioles with several individual to almost all pos EC; vAg covering luminal surface; rare adjacent alveoli with pos type I and II pc  **BLN**: NE |  |  |  |
| **Infected, Nafamostat**  **4 mg/kg**  [3.1]  Fig 3B | **2** | **Lung (HE)**: rare degen BEC | 15.82 | 12.3 | 18.47 |
|  |  | **Nose**: patches of pos EC in entire nasal mucosa (REC and OEC)  **Trachea**: rare individual pos EC  **Lung**: many bronchioles with individual to stretches of pos, partly degen EC; some vAg covering luminal surface; adjacent focal area of alveoli with pos type I and II pc  **BLN**: NE |  |  |  |
| **Infected,**  **Nafamostat 4 mg/kg**  [3.2] | **2** | **Lung (HE)**: rare degen BEC | 1 | 1.73 | 3.88 |
|  |  | **Nose**: patches of pos REC and OEC  **Trachea**: small amount of vAg attached to EC  **Lung**: several bronchioles with individual to a few pos, partly degen EC; some vAg covering luminal surface  **BLN**: NE |  |  |  |
| **Infected,**  **Nafamostat 4 mg/kg**  [3.3]  Fig 3A | **2** | **Lung (HE)**: NHA | 32.53 | 23,43 | 29.8 |
|  |  | **Nose**: patches of pos REC and OEC  **Trachea**: small amount of vAg attached to EC  **Lung**: a few small patches of alveoli with pos type I and II pc  **BLN**: neg |  |  |  |
| **Infected, Nafamostat 4 mg/kg**  [3.4] | **2** | **Lung (HE)**: NHA | 5.63 | 6.07 | 11.37 |
|  |  | **Nose**: patches of several to numerous pos intact OEC, occ pos cells in underlying BG and in nerve bundles  **Trachea**: neg  **Lung**: a few pos type I and II pc  **BLN**: neg |  |  |  |
| **Infected,**  **Nafamostat 4 mg/kg;**  **Apilimod 1 mg/kg**  [4.1] | **2** | **Lung (HE)**: NHA | 1.79 | 2.1 | 2.92 |
|  |  | **Nose**: patches of pos REC and OEC, focal pos cells in underlying BG and in nerve bundles  **Trachea**: neg  **Lung**: one small patch of a few pos type I and II pc  **BLN**: neg |  |  |  |
| **Infected,**  **Nafamostat 4 mg/kg;**  **Apilimod 1 mg/kg**  [4.2] | **2** | **Lung (HE)**: NHA | 0 | 0.02 | 0.22 |
|  |  | **Nose**: patches of pos REC (apical and dorsal) and OEC, focus of pos cells in underlying BG and in nerve bundles  **Trachea**: neg  **Lung**: two small patches of a few pos type I and II pc  **BLN**: neg |  |  |  |
| **Infected,**  **Nafamostat 4 mg/kg;**  **Apilimod 1 mg/kg**  [4.3] | **2** | **Lung (HE)**: small focal subpleural area with activated type II pc and some infiltrating NL | 0 | 0.03 | 0.11 |
|  |  | **Nose**: patches of pos REC and OEC  **Trachea**: neg  **Lung**: small focal subpleural area with pos type II pc (see HE)  **BLN**: neg |  |  |  |
| **Infected,**  **Nafamostat 4 mg/kg;**  **Apilimod 1 mg/kg**  [4.4] | **2** | **Lung (HE)**: small focal subpleural area with activated type II pc and some infiltrating NL | 2.3 | 1.58 | 2.6 |
|  |  | **Nose**: patches of pos REC and OEC  **Trachea**: neg  **Lung**: two small focal subpleural areas with pos type I and II pc (see also HE)  **BLN**: neg |  |  |  |
| **Infected,**  **Nafamostat 4 mg/kg;**  **Apilimod 2 mg/kg**  [5.1]  Fig 3A | **2** | **Lung (HE)**: NHA | 0 | 0.09 | 0.28 |
|  |  | **Nose**: small patches of pos intact OEC adjacent to olfactory bulb  **Trachea**: neg  **Lung**: one focal subpleural area with pos type I and II pc  **BLN**: neg |  |  |  |
| **Infected,**  **Nafamostat 4 mg/kg;**  **Apilimod 2 mg/kg**  [5.2] | **2** | **Lung (HE)**: NHA | 0.65 | 0.54 | 1.41 |
|  |  | **Nose**: small patches of pos intact OEC adjacent to olfactory bulb  **Trachea**: neg  **Lung**: one focal subpleural area with pos type I and II pc  **BLN**: NE |  |  |  |
| **Infected,**  **Nafamostat 4 mg/kg;**  **Apilimod 2 mg/kg**  [5.3] | **2** | **Lung (HE)**: NHA | 0 | 0 | 0 |
|  |  | **Nose**: small patches of pos intact OEC adjacent to olfactory bulb  **Trachea**: neg  **Lung**: neg  **BLN**: NE |  |  |  |
| **Infected,**  **Nafamostat 4 mg/kg;**  **Apilimod 2 mg/kg**  [5.4]  Fig 3B | **2** | **Lung (HE)**: a few small, LC-dominated pb leukocyte aggregates | 0 | 0 | 0 |
|  |  | **Nose**: large patches of pos intact OEC adjacent to olfactory bulb, a few pos cells in underlying BG and in nerve bundles  **Trachea**: neg  **Lung**: neg  **BLN**: NE |  |  |  |
| **Experiment 2** | | | | | |
| **Infected, not treated**  [1.1]  Fig 4B | **2** | **Lung (HE)**: several bronchioles with degen and loss of EC; mild pb mononuclear infiltration; focal areas with activated type II pc, some degen AEC and infiltrating NL; mild vasculitis and pv infiltration | 51.05 | 61.99 | 111.73 |
|  |  | **Nose**: abundant pos REC (apical), occ pos OEC  **Trachea**: rare intact pos EC, pos debris in lumen  **Lung**: several bronchioles with very abundant pos EC; vAg in debris in lumen; disseminated pos type I and II pc (individual, small groups)  **BLN**: several pos macrophages/DC |  |  |  |
| **Infected, not treated**  [1.2]  Fig 4A | **2** | **Lung (HE)**: several bronchioles with degen and loss of EC; mild pb mononuclear infiltration; focal areas with activated type II pc, some degen AEC and infiltrating NL; mild vasculitis and pv infiltration | 527.80 | 686.85 | 128.34 |
|  |  | **Nose**: abundant pos OEC (mid to caudal nasal mucosa)  **Trachea**: rare intact pos EC, pos debris in lumen  **Lung**: several bronchioles with very abundant pos EC; vAg in debris in lumen; disseminated pos type I and II pc (individual, small groups)  **BLN**: several pos macrophages/DC |  |  |  |
| **Infected, not treated**  [1.3] | **2** | **Lung (HE)**: several bronchioles with degen and loss of EC; mild pb mononuclear infiltration; focal areas with activated type II pc, some degen AEC and infiltrating NL; mild vasculitis and pv infiltration | 37.11 | 23.49 | 69.74 |
|  |  | **Nose**: abundant pos REC and OEC (entire nasal mucosa)  **Trachea**: pos debris in lumen  **Lung**: several bronchioles with very abundant pos EC; vAg in debris in lumen; disseminated pos type I and II pc (individual, small groups)  **BLN**: several pos macrophages/DC |  |  |  |
| **Infected,**  **Apilimod 2 mg/kg**  [2.1] | **2** | **Lung (HE)**: several bronchioles with degen and loss of EC; mild pb mononuclear infiltration; focal areas with activated type II pc, some degen AEC and infiltrating NL; mild vasculitis and pv infiltration | 186.61 | 292.82 | 118.92 |
|  |  | **Nose**: patches of pos REC and OEC (entire nasal mucosa)  **Trachea**: patches of pos EC, pos debris in lumen  **Lung**: several bronchioles with very abundant pos EC; vAg in debris in lumen; some alveoli with individual to several pos type I and II pc  **BLN**: NE |  |  |  |
| **Infected,**  **Apilimod 2 mg/kg**  [2.2]  Fig 3A | **2** | **Lung (HE)**: several bronchioles with degen and loss of EC; mild pb mononuclear infiltration; focal areas with activated type II pc, some degen AEC and infiltrating NL; mild vasculitis and pv infiltration | 606.29 | 1139.24 | 450.02 |
|  |  | **Nose**: patches of pos REC and OEC (entire nasal mucosa)  **Trachea**: several large patches of pos EC, pos debris in lumen  **Lung**: several bronchioles with very abundant pos EC; vAg in debris in lumen; some alveoli with individual to several pos type I and II pc  **BLN**: NE |  |  |  |
| **Infected,**  **Apilimod 2 mg/kg**  [2.3]  Fig 3B | **2** | **Lung (HE)**: several bronchioles with degen and loss of EC; mild pb mononuclear infiltration; focal areas with activated type II pc, some degen AEC and infiltrating NL; mild vasculitis and pv infiltration | 40.33 | 78.46 | 239.50 |
|  |  | **Nose**: patches of pos REC and OEC (mid to caudal nasal mucosa)  **Trachea**: several large patches of pos EC, pos debris in lumen  **Lung**: several bronchioles with very abundant pos EC; vAg in debris in lumen; some alveoli with individual to several pos type I and II pc  **BLN**: a few pos macrophages/DC |  |  |  |
| **Infected,**  **Nafamostat 0.8 mg/kg;**  **Apilimod 0.4 mg/kg**  [3.1] | **2** | **Lung (HE)**: NHA | 0 | 0 | 0.02 |
|  |  | **Nose**: neg  **Trachea**: neg  **Lung**: neg  **BLN**: neg |  |  |  |
| **Infected,**  **Nafamostat 0.8 mg/kg;**  **Apilimod 0.4 mg/kg**  [3.2] | **2** | **Lung (HE)**: NHA | 0 | 0 | 0.02 |
|  |  | **Nose**: large area with large patches of pos intact OEC (dorsocaudal nasal mucosa)  **Trachea**: neg  **Lung**: neg  **BLN**: neg |  |  |  |
| **Infected,**  **Nafamostat 0.8 mg/kg;**  **Apilimod 0.4 mg/kg**  [3.3] | **2** | **Lung (HE)**: NHA | 0 | 0 | 0.01 |
|  |  | **Nose**: a few individual pos intact OEC (dorsocaudal nasal mucosa)  **Trachea**: neg  **Lung**: neg  **BLN**: neg |  |  |  |
| **Infected,**  **Nafamostat 0.8 mg/kg;**  **Apilimod 0.4 mg/kg**  [3.4] | **2** | **Lung (HE)**: NHA | 0.06 | 0.05 | 0.12 |
|  |  | **Nose**: neg  **Trachea**: neg  **Lung**: neg  **BLN**: neg |  |  |  |
| **Infected,**  **Nafamostat 0.4 mg/kg;**  **Apilimod 0.4 mg/kg**  [4.1] | **2** | **Lung (HE)**: NHA | 0 | 0.11 | 0.20 |
|  |  | **Nose**: a few individual and patches of pos intact OEC (caudal nasal mucosa)  **Trachea**: neg  **Lung**: neg  **BLN**: neg |  |  |  |
| **Infected,**  **Nafamostat 0.4 mg/kg;**  **Apilimod 0.4 mg/kg**  [4.2] | **2** | **Lung (HE)**: scattered individual degen BEC | 8.13 | 6.16 | 6.00 |
|  |  | **Nose**: a few individual and patches of pos intact REC (apical nasal mucosa) and OEC (caudal nasal mucosa)  **Trachea**: neg  **Lung**: a few patches of pos, partly degen BEC and rare type I and II pc  **BLN**: neg |  |  |  |
| **Infected,**  **Nafamostat 0.4 mg/kg;**  **Apilimod 0.4 mg/kg**  [4.3] | **2** | **Lung (HE)**: small focal subpleural area with activated type II pc and some infiltrating NL | 6.12 | 7.18 | 8.96 |
|  |  | **Nose**: abundant pos REC and OEC (apical to mid nasal mucosa)  **Trachea**: neg  **Lung**: several patches of alveoli with pos type II pc  **BLN**: a few pos macrophages/DC |  |  |  |
| **Infected,**  **Nafamostat 0.4 mg/kg;**  **Apilimod 0.4 mg/kg**  [4.4] | **2** | **Lung (HE)**: NHA | 0 | 0 | 0.05 |
|  |  | **Nose**: abundant pos REC and OEC (entire nasal mucosa)  **Trachea**: neg  **Lung**: neg  **BLN**: NE |  |  |  |
| **Infected,**  **Nafamostat 4 mg/kg;**  **Apilimod 2mg/kg;**  **from 6 hpi**  [5.1]  Fig 4B | **2** | **Lung (HE)**: NHA | 59.87 | 72.70 | 124.83 |
|  |  | **Nose**: individual and patches of pos intact REC and OEC (entire nasal mucosa)  **Trachea**: rare pos EC  **Lung**: several bronchioles with few to many pos, occ degen BEC, occ adjacent pos type II pc  **BLN**: neg |  |  |  |
| **Infected,**  **Nafamostat 4 mg/kg;**  **Apilimod 2mg/kg;**  **from 6 hpi**  [5.2] | **2** | **Lung (HE)**: scattered individual degen BEC | 102.81 | 127.46 | 258.47 |
|  |  | **Nose**: abundant pos intact REC and OEC entire nasal mucosa)  **Trachea**: rare pos EC  **Lung**: several bronchioles with some to many pos, occ degen BEC, occ adjacent pos type II pc  **BLN**: numerous pos macrophages/DC |  |  |  |
| **Infected,**  **Nafamostat 4 mg/kg;**  **Apilimod 2mg/kg;**  **from 6 hpi**  [5.3] | **2** | **Lung (HE)**: scattered individual degen BEC | 411.25 | 509.82 | 244.53 |
|  |  | **Nose**: abundant pos intact REC and OEC (entire nasal mucosa)  **Trachea**: NE  **Lung**: most bronchioles with some to many pos, occ degen BEC, occ adjacent pos type II pc  **BLN**: numerous pos macrophages/DC |  |  |  |
| **Infected,**  **Nafamostat 4 mg/kg;**  **Apilimod 2mg/kg;**  **from 6 hpi**  [5.4] | **2** | **Lung (HE)**: scattered individual degenerate BEC | 38.69 | 62.85 | 197.25 |
|  |  | **Nose**: abundant pos intact REC and OEC (entire nasal mucosa)  **Trachea**: rare pos EC  **Lung**: several bronchioles with some to many pos, occ degen EC, occ adjacent pos type II pc  **BLN**: a few pos macrophages/DC |  |  |  |
| **Experiment 3** | | | | | |
| **Infected, not treated**  [1.1] | **2** | **Lung (HE)**: several bronchioles with degen and loss of EC; mild pb mononuclear infiltration; focal areas with activated type II pc | 231.87 | 158.01 |  |
|  |  | **Nose**: abundant pos intact REC, some pos OEC (entire nasal mucosa)  **Trachea**: a few pos EC, some pos debris in lumen  **Lung**: numerous bronchioles with several to abundant pos EC, pos debris in lumen; disseminated alveoli with individual to several pos type I and II pc  **BLN**: several pos macrophages/DC |  |  |  |
| **Infected, not treated**  [1.2] | **2** | **Lung (HE)**: several bronchioles with degen and loss of EC; mild pb mononuclear infiltration; focal areas with activated type II pc | 65.67 | 149.48 |  |
|  |  | **Nose**: abundant pos intact REC (apical and mid nasal mucosa), some pos OEC  **Trachea**: very rare pos EC, some pos debris in lumen  **Lung**: numerous bronchioles with several to abundant pos EC, pos debris in lumen; disseminated alveoli with individual to several pos type I and II pc  **BLN**: numerous pos macrophages/DC |  |  |  |
| **Infected, not treated**  [1.3]  Fig 5C | **2** | **Lung (HE)**: several bronchioles with degen and loss of EC; mild pb mononuclear infiltration; focal areas with activated type II pc | 65.67 | 42.34 |  |
|  |  | **Nose**: abundant pos intact REC (apical and mid nasal mucosa), some pos OEC  **Trachea**: NE  **Lung**: numerous bronchioles with several to abundant pos EC, pos debris in lumen; disseminated alveoli with individual to several pos type I and II pc  **BLN**: NE |  |  |  |
| **Infected, not treated**  [2.1] | **4** | **Lung (HE)**: NHA | 0 | 0 |  |
|  |  | **Nose**: a few intact pos OEC (caudal nasal mucosa)  **Trachea**: neg  **Lung**: neg  **BLN**: neg |  |  |  |
| **Infected, not treated**  [2.2]  Fig 5C | **4** | **Lung**: mild vasculitis and pv LC infiltration | 0 | 0 |  |
|  |  | **Nose**: several individual intact pos OEC (caudal nasal mucosa)  **Trachea**: neg  **Lung**: multifocal individual pos type I amd II pc and macrophages  **BLN**: neg |  |  |  |
| **Infected,**  **Nafamostat 4 mg/kg;**  **Apilimod 2mg/kg**  [3.1] | **2** | **Lung (HE)**: very mild pv and pb LC infiltration | 0 | 0 |  |
|  |  | **Nose**: a few intact pos OEC (caudal nasal mucosa)  **Trachea**: neg  **Lung**: neg  **BLN**: NE |  |  |  |
| **Infected,**  **Nafamostat 4 mg/kg;**  **Apilimod 2mg/kg**  [3.2]  Fig 5C | **2** | **Lung (HE)**: NHA | 0 | 0 |  |
|  |  | **Nose**: neg  **Trachea**: neg  **Lung**: neg  **BLN**: NE |  |  |  |
| **Infected,**  **Nafamostat 4 mg/kg;**  **Apilimod 2mg/kg**  [3.3] | **2** | **Lung (HE)**: NHA | 0 | 0 |  |
|  |  | **Nose**: neg  **Trachea**: neg  **Lung**: neg  **BLN**: NE |  |  |  |
| **Infected,**  **Nafamostat 4 mg/kg;**  **Apilimod 2mg/kg**  [3.4] | **2** | **Lung (HE)**: NHA | 0 | 0 |  |
|  |  | **Nose**: abundant pos REC and OEC (mainly apical and mid nasal mucosa)  **Trachea**: neg  **Lung**: neg  **BLN**: neg |  |  |  |
| **Infected,**  **Nafamostat 4 mg/kg;**  **Apilimod 2mg/kg;**  **for 2 days**  [4.1]  Fig 5C | **4** | **Lung (HE)**: mild multifocal areas with activated type II PC; focal area with some desquamed type II pc/alv macrophages; mild focal pv LC infiltration | 0 | 0 |  |
|  |  | **Nose**: a few intact REC (apical nasal mucosa)  **Trachea**: neg  **Lung**: a few pos type II pc/alv macrophages in affected area (see HE)  **BLN**: very few pos macrophages |  |  |  |
| **Infected,**  **Nafamostat 4 mg/kg;**  **Apilimod 2mg/kg;**  **for 2 days**  [4.2] | **4** | **Lung (HE)**: NHA | 0 | 0 |  |
|  |  | **Nose**: neg  **Trachea**: neg  **Lung**: a few small groups of pos BEC  **BLN**: very few pos macrophages |  |  |  |
| **Infected,**  **Nafamostat 4 mg/kg;**  **Apilimod 2mg/kg;**  **for 2 days**  [4.3] | **4** | **Lung (HE)**: NHA | 0 | 0 |  |
|  |  | **Nose**: neg  **Trachea**: neg  **Lung**: neg  **BLN**: neg |  |  |  |
| **Infected,**  **Nafamostat 4 mg/kg;**  **Apilimod 2mg/kg;**  **for 2 days**  [4.4] | **4** | **Lung (HE)**: multifocal areas with activated type II PC; a few focal areas with some desquamed type II pc/alv macrophages; mild focal pv LC infiltration | 0 | 0.01 |  |
|  |  | **Nose**: a few intact pos OEC (caudal nasal mucosa)  **Trachea**: neg  **Lung**: rare bronchioles with a few pos EC; a few pos type II pc/alv macrophages in affected area (see HE)  **BLN**: neg |  |  |  |
| **Infected,**  **Nafamostat 4 mg/kg;**  **Apilimod 2mg/kg;**  **from 3 hpi**  [5.1] | **2** | **Lung (HE)**: focal areas with activated type II pc and occ syncytial cells, degen cells, NL and desquamed type II pc/alv macrophages; degen cells in bronchiolar lumina; mild vasculitis | 0.15 | 45.38 |  |
|  |  | **Nose**: abundant pos OEC (mainly mid and caudal nasal mucosa)  **Trachea**: neg  **Lung**: many bronchioles with individual to abundant pos EC; some (adjacent) pos type II pc and macrophages; focal areas (see HE) with pos AEC and macrophages, some pos degen EC in bronchiolar lumina  **BLN**: NE |  |  |  |
| **Infected,**  **Nafamostat**  **4 mg/kg;**  **Apilimod 2mg/kg from 3 hpi**  [5.2] | **2** | **Lung (HE)**: multifocal activated type II pc | 99.54 | 100.70 |  |
|  |  | **Nose**: abundant pos OEC (mid and caudal nasal mucosa)  **Trachea**: neg  **Lung**: several bronchioles with some to abundant pos EC; patches of alveoli with pos type II and fewer type I pc  **BLN**: NE |  |  |  |
| **Infected,**  **Nafamostat 4 mg/kg;**  **Apilimod 2mg/kg;**  **from 3 hpi**  [5.3]  Fig. 4A | **2** | **Lung (HE)**: focal areas with activated type II pc and occ degen cells, NL and desquamed type II pc/alv macrophages; a few degen cells in bronchiolar lumina | 117.55 | 147.43 |  |
|  |  | **Nose**: moderate number of pos OEC (mid and caudal nasal mucosa)  **Trachea**: neg  **Lung**: several bronchioles with individual to numerous pos EC and some desquamed pos EC in lumen; vAg covering luminal surface; patches of alveoli with pos type II and fewer type I pc  **BLN**: NE |  |  |  |
| **Infected,**  **Nafamostat 4 mg/kg;**  **Apilimod 2mg/kg;**  **from 3 hpi**  [5.4] | **2** | **Lung (HE)**: focal areas with activated type II pc and occ degen cells, NL and desquamed type II pc/alv macrophages; one bronchiole with a few degen cells in lumen | 442.00 | 348.22 |  |
|  |  | **Nose**: moderate number of pos OEC (mid and caudal nasal mucosa)  **Trachea**: neg  **Lung**: many bronchioles with individual to numerous pos EC and some desquamed pos EC in lumen; vAg covering luminal surface; patches of alveoli with pos type II and fewer type I pc; focal areas (see HE) with pos AEC and macrophages  **BLN**: neg |  |  |  |

**Legend**: alv – alveolar; BEC – bronchiolar epithelial cells; BG – Bowman’s glands; BLN – bronchial lymph node; DC – dendritic cells; degen – degenerate; EC – epithelial cells; F – female; HE – histological features assessed in a hematoxylin-eosin stained section; LC – lymphocyte; M – male; NB – nerve bundles; NE – not examined; neg – negative; NHA – no histological abnormality; NL – neutrophilic leukocytes; nose – nasal mucosa including the respiratory and olfactory epithelium; occ – occasional; OEC – olfactory epithelial cells; pb – peribronchiolar; pc – pneumocytes; pos – positive; pv – perivascular; REC – respiratory epithelial cells; vAg – viral antigen;

*Head: This implies examination of the two halves of the head after longitudinal sawing in the midline, allowing examination of the nasal mucosa and the brain including the olfactory bulb. Since there was no evidence of viral antigen expression in brain including olfacory bulb of any animal, this is not reported for each animal in the table.
